# PlanktonFlow : hands-on deep-learning classification of plankton images for biologists

**DOI:** 10.1101/2025.09.19.677346

**Authors:** Hugo Walter, Caroline Gorzerino, Marc Collinet, Béatrice Porcon, François Martignac, Eric Edeline

## Abstract

High throughput image-acquisition devices tremendously increase our capacity to observe biodiversity. However, for many biologists, the high-performance deep learning models that are needed to make biological sense out of very-large image sets remain difficult to implement.

To fill this gap in biologists’ toolkit, we developed PlanktonFlow, a Python pipeline that streamlines the automation of plankton-image taxonomic assignment. PlanktonFlow makes it easy for inexperienced users to run a whole sequence of (i) automated image pre-processing and augmentation of rare classes, (ii) training up to four different high-performance convolution neural networks (CNNs: ResNet, DenseNet, EfficientNet, and YOLO), (iii) computing model classification-performance metrics so as to choose the best-performing model, and (iv) running inference on novel image sets. PlanktonFlow further includes routines to easily fine tune model hyper-parameters and optimize model’s performances.

Using a tutorial style, we demonstrate the usage of PlanktonFlow to analyse freshwater-plankton images produced with the FlowCAM, comparing the relative classification performances of the four optimized CNN architectures. For a baseline comparison with a reference tool used by plankton biologists, we further assessed the classification performances of the EcoTaxa web-service when used without any eye validation in a pure-prediction mode. In line with a previous study on a benchmark plankton dataset, we found that EfficientNet-B5 achieved the highest macro-averaged F1 Score, outperforming other CNN models, which all surpassed EcoTaxa. Hyper-parameter optimization was key to improving model performances.

To ease an appropriation and further developments by the community, PlanktonFlow is open source, comes with a detailed documentation, and has a modular structure. We foresee that future work could integrate new deep-learning architectures (e.g., vision transformers, semi-supervised learning), and test the pipeline on images produced by other devices or from other taxonomic groups.

## Introduction

High-throughput biodiversity monitoring, as enabled by modern automatic sensors, provides us with an unprecedented capacity to track rich, multidimensional data streams and, ultimately, to accurately understand and forecast ecological dynamics across broad spatial and temporal scales (Besson et al., 2022, Weinstein, 2018). In aquatic systems, a diversity of imaging-in-flow and camera systems now capture a wide spectrum of plankton sizes and taxonomic diversity. Submersible, high-frequency instruments like the Imaging FlowCytobot (Olson and Sosik, 2007), the Dual Scripps Plankton Camera (DSPC, Merz et al., 2021) or the Underwater Vision Profiler (UVP, Picheral et al., 2010) operate autonomously *in situ* to capture continuous records of plankton. In parallel, benchtop imaging flow cytometers for plankton, such as the FlowCAM^®^ (Sieracki et al., 1998) or the PlanktoScope (Pollina et al., 2022), are widely used to analyse water samples *ex-situ* in the laboratory or under shipboard settings.

However, the volume and heterogeneity of images produced by these machines often exceed the routine-processing capacity of traditional analysis workflows, and thus create bottlenecks that hinder scientific progress. Softwares shipped with commercial instruments (for example, FlowCAM’s VisualSpreadsheet) typically require substantial manual curation and make it impossible to process large datasets (Owen et al., 2022). To scale up classification, the Laboratoire d’Océanographie de Villefranche developed EcoTaxa, a web-based platform intended for computer-assisted image classification, which is now the standard tool used by biologists for plankton analysis (Irisson et al., 2022; Picheral et al., 2025).

EcoTaxa provides a user-friendly collaborative graphical interface. Based on a validated dataset, EcoTaxa predicts an object class using random forests fed with multiple image features, including convolution neural networks (CNNs)-extracted features (see Figure S1, Walter et al., 2025). Predicted classes are then intended to be eye-validated. However, when projects are so large that they surpass human-validation capacities, one may use EcoTaxa without any further eye validation in a “pure-prediction mode”, which consists of either using directly the classification provided by random forests, or of retaining only images with the highest random-forest prediction score (Faillettaz et al., 2016). Unfortunately however, EcoTaxa lacks prediction-quality metrics (accuracy, recall, confusion matrix…), making it difficult to assess its performances in a pure-prediction mode.

At the same time, recent studies show on various benchmark plankton datasets that modern deep-learning approaches (CNNs, Vision Transformers) can achieve classification accuracies that potentially surpass those achieved by EcoTaxa (Kyathanahally et al., 2021, Kyathanahally et al., 2022, Eerola et al., 2024). However, several obstacles prevent the adoption of these novel methods by EcoTaxa users. The codes and data used to run novel methods are sometimes publicly available, but these scripts may run proprietary software (Lumini and Nanni, 2019), and/or their documentation may be too limited for beginners to be able to adapt the scripts to their own needs (e.g., Kyathanahally et al., 2021, Kraft et al., 2022, Kyathanahally et al., 2022). In fact, the computational/engineering overhead required to train, deploy and maintain robust models in operational settings are often beyond-of-reach for non-specialists. Another hindrance to the reutilization of published scripts is their frequent lack of routines for image pre-treatment (e.g., augmentation of rare classes) or performance measurements (but see Maracani et al., 2023). Yet, performance comparison among architectures is crucial because variation among datasets often precludes a direct transfer of methods among instruments and environments, a problem known as “dataset shifts” (Chen et al., 2025), and also because severe class imbalance due to rare taxa imposes custom-trained models. For all these reasons, modern deep-learning methods remain beyond of reach to many biologists.

To fill this gap in biologists’ toolkit, we developed PlanktonFlow, a novel pipeline that streamlines deep-learning annalysis of large plankton image sets. The pipeline ships with transparent, reproducible training recipes, clear step-by-step instructions, and all of the configuration files needed to adapt the pipeline to other instruments or taxa. From an already-built dataset of manually-labelled images, PlanktonFlow was designed to make it easy for beginners to:

- pre-treat their labelled images for scale-bar removel (optional),
- augment the number of images for rare classes in their dataset (optional),
- train up to four different CNNs on their dataset and compare their performances. Specifically, users can choose either pretrained CNNs ready for immediate use, or to automatically train a CNN on their own dataset, choosing among ResNet, DenseNet, EfficientNet, and YOLO architectures. PlanktonFlow further allows for hyperparameter tuning of the enclosed CNNs.
- Extensively evaluate and compare models classification performances.
- Run predictions from the best-performing model on a novel dataset.

To demonstrate the usage of PlanktonFlow, we here implement the whole pipeline on a dataset of labelled plankton images from our own freshwater mesocosm experiments. Images of ethanol-fixed samples were produced using the FlowCAM, and were manually-labelled using ExoTaxa and a single eye validation. We deliberately chose not to work with a benchmark dataset, because benchmarks are less likely to represent the whole diversity of object classes encountered under operational conditions. Compared to benchmark datasets, our custom dataset was maybe more difficult to classifify, which would decrease the absolute performances of classifiers, but would not influence their relative performances.

As a baseline for performance comparison, we use EcoTaxa in a pure-prediction mode, so as to test for an expected added value of modern classifiers compared to a standard classification tool used by plankton biologists.

## Material and methods

### Dataset

#### Raw dataset

The plankton image dataset used in this study was acquired from a long-term mesocosm experiment on the effects of invasive crayfish on freshwater ecosystems (Dézerald et al., 2023). 12 outdoor, fishless mesocosms were installed in 2020 at the PEARL, the INRAE experimental platform for aquatic ecology and ecotoxicology (U3E) located in Rennes, France (Platform PEARL). The mesocosms were initially seeded with pond sediments, such that a rich vegetal and animal community rapidly developed. Organisms were also free to colonise mesocosms by aerial dispersal. In 2021, red swamp crayfish (*Procambarus clarkii*) were introduced in eight mesocosms and soon eradicated all the vegetation. Monthly water samples were taken from the 12 mesocosms in 2022 and 2023 (24 months). Hence, our image dataset contained diverse plankton taxa from both vegetated and non-vegetated natural freshwater ecosystems.

Imaging was performed using the FlowCAM (Fluid Imaging Technologies Inc.), a high-throughput imaging cytometer designed for particle analysis in liquid media. In image-based studies, the taxonomic resolution attainable by either humans or deep-learning methods critically depends on image resolution. To increase the image resolution of our dataset, we used two separate Flow-CAM instruments optimized for different organismal size ranges: a FlowCAM 8110 dedicated to microplankton (50 to 200 *µm*) and a FlowCAM Macro dedicated to macroplankton (0.2 to 2*mm*). High-resolution images in .tiff format produced by the FlowCAMs were segmented using versions 8.22 to 8.26 of the ZooProcess ImageJ macro (Jalabert et al., 2025), that returned low-resolution images (approximately 400 by 400 *px* ), each containing one object. The objects exhibited a wide spectrum of visual complexity, including variations in pose, orientation, occlusion, transparency, and morphological deformation. This heterogeneity reflected the natural complexity of water samples including target planktonic organisms, non-target biological material (e.g., terrestrial organisms), and detritus and other particles.

The low-resolution images produced by ZooProcess are the raw material used to build the dataset and to feed PlanktonFlow. Images were uploaded to EcoTaxa for 100% manual construction of our dataset of labelled images (i.e., none of the models assessed below previously “saw” the data). In building the dataset, we retained a large number of classes reflecting the whole diversity of objects effectively encountered, rather than reducing the label space to ease subsequent automatic classification. We also chose not to double-check the dataset for labelling errors (a highly time-consuming task), because our goal was *not* to establish a new benchmark reference model or benchmark model comparison, but rather to develop an operational pipeline. Therefore, our dataset is not a reference dataset like is, for instance, WHOI-Plankton (Orenstein et al., 2015).

#### Data pre-processing

To ensure the reliability and statistical relevancy of the training data, we applied a pruning strategy that removed extremely rare classes. The dataset exhibited a strongly skewed class distribution, which is typical of ecological samples: the top 33 most abundant taxa accounted for over 70% of all annotated images, while the remaining 73 taxa collectively contributed the remaining 30%. Specifically, any class with fewer than 100 manually-labelled images was excluded from the final dataset. This decision was motivated by two factors: (1) very low sample sizes tend to introduce high-variance gradients that hinder stable model convergence, and (2) such elusive organisms are rarely meaningful ecologically and often will not be used in later intended research. However, quantifying rare taxa using deep learning is possible, and simply requires to label more images for these taxa (here a suggested N=100), provided that enough images are available in the whole image set.

After pruning, the number of valid classes was reduced from *N*_*original*_=106 to *N*_*final*_=76, with the removed classes contributing marginally to the overall image volume (Figure 1). Among the 76 classes, 51 were taxonomic classes including six species (e.g., *Brachionus calyciflorus*), 28 genera (e.g., *Chydorus*), seven families (e.g., Daphniidae), one infra-class (Oligochaeta), four classes (e.g., Copepoda), one infra-order (Cladocera), two orders (e.g., Cyclopoida), and two Phyla (e.g., Rotifera) (see Figure S2 in Walter et al., 2025).

**Figure 1.**
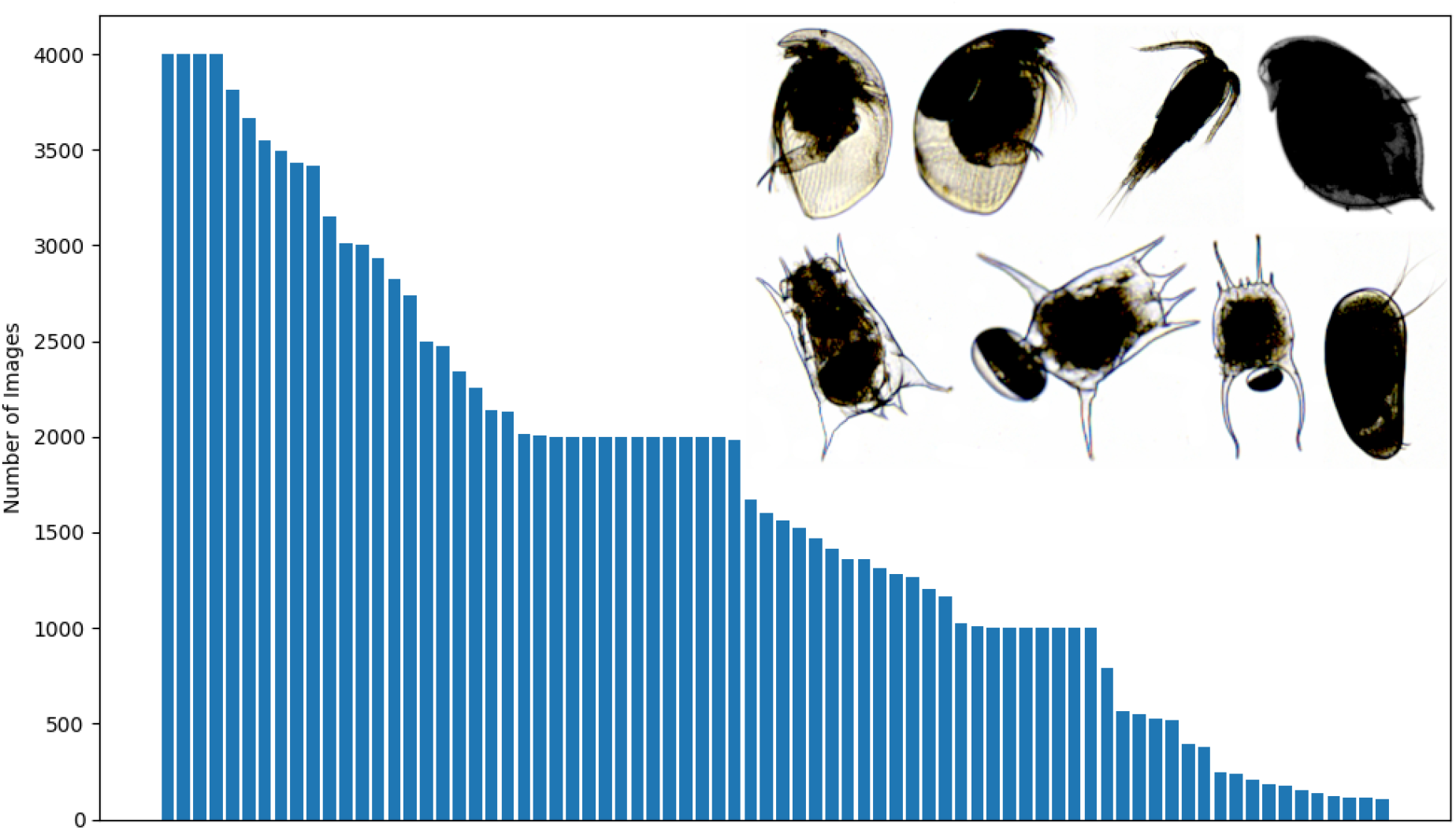
Raw class distribution after pruning rare classes and before data augmentation. The inset shows sample images of zooplankton from freshwater mesocosms.

A critical preprocessing step in the pipeline involved detecting and removing scale bars that were embedded in some of the original images. Inconsistency in the presence of scale bars was due to varying settings in our use of the different versions of ZooProcess. To prevent data leakage due to the model learning scale bars rather than biological features, we implemented an automated pipeline to remove scale bars. Initial tests using traditional computer vision techniques (template matching) proved unreliable due to slight inter-image variability in the scale bar’s pixel rendering and contrast. We therefore fine-tuned a YOLOv8 object detection model for this task. On a held-out test set (N=160), the model achieved 100% precision and recall, with zero false positives or false negatives. Since scale bars in our images did not overlap with the plankton objects, this perfect detection rate ensured that all artifacts were completely removed without compromising biological data.

Finally, the pre-processed image set was split into three disjoint subsets in a stratified manner across the 76 classes, ensuring that rare classes remained proportionally represented similarly across all subsets: 70% for training (90,723 images), 20% for validation (25,921 images), and 10% for testing (12,961 images).

#### Data Augmentation

The performances of deep learning models often deteriorate when exposed to imbalanced training distributions, especially in multi-class settings with sparse and heterogeneous categories. In the context of our plankton classification task, pronounced class imbalance was observed across taxa, primarily due to natural ecological abundance gradients. To mitigate this issue and to enhance the model’s generalization capability, we implemented a *class-aware data augmentation strategy* aimed at selectively enriching under-represented categories in the training dataset.

Specifically, we augmented data using the Albumentations library (Buslaev et al., 2018), where each transformation is applied independently with a given probability. The transformations included horizontal flipping (50%), vertical flipping (20%), 90-degree rotation (30%), brightness and contrast adjustment (40%), and hue/saturation modulation (30%). Since these operations are sampled separately, multiple augmentations may be applied to the same image.

To standardise the training distribution, we defined an *augmentation threshold* of 1,500 images per class. For any class below this threshold, we calculated the required number of synthetic samples to produce, and we distributed the augmentation load across the available original images using a capped multiplier (maximum 10 augmentations per original). This process ensured a controlled and balanced enrichment of the dataset while maintaining fidelity to the original morphological patterns.

The class-aware data augmentation strategy was applied exclusively to the training set. As a result, the final dataset partitions used for model training and evaluation were as follows:

- *D*_train_ : Training Set: 159,687 images (original + augmented),
- *D*_val_ : Validation Set: 25,921 images (original only),
- *D*_test_ : Test Set: 12,961 images (original only).

### Model architectures

The problem of dataset shifts makes it crucial to evaluate different deep-learning architectures on the dataset at hand (Chen et al., 2025). To illustrate PlanktonFlow’s capabilities on this task, we compared the classification performances of four different CNN models, all of which are enclosed in PlanktonFlow: ResNet (He et al., 2016), DenseNet (Huang et al., 2018), EfficientNet (Tan and Le, 2020) and YOLOv11 (Khanam and Hussain, 2024).

We chose ResNet in our candidate-model set for its use of residual connections, which facilitates gradient flow and enables the training of deep networks without degradation. Variants like ResNet-50 offer a solid balance between accuracy and resource use, making them a common baseline for transfer learning. DenseNet connects each layer to every other layer in a feed-forward manner, improving gradient propagation and promoting feature reuse. This architecture achieves strong generalization with fewer parameters, making it well-suited for complex but data-limited tasks. EfficientNet applies compound scaling to jointly optimize network depth, width, and input resolution, achieving high accuracy with fewer parameters and FLOPs (Floating Point Operations, measure computational complexity). Its scalable variants (B0–B7) offer flexibility between speed and performance, with smaller models enabling fast prototyping. Finally, YOLO is a multi-task CNN that is advertised by its developers to excel at object detection, tracking, instance segmentation, image classification, and pose estimation tasks^1^. We included YOLOV11 in our panel of CNNs because it is arguably the most user-friendly deep learning workflow, and as such it is easily accessible to beginners.

For a baseline comparison with the standard tool used by many plankton biologists, we further measured the classification performances provided by EcoTaxa in a pure-prediction mode withtout any further human validation. All four deep-learning architectures were subject to the exact same training, validation and testing procedures with *D*_train_, *D*_val_ and *D*_test_, respectively (see Supplementary Information, Walter et al., 2025). In contast, because the platform makes it impossible to have a validation step, EcoTaxa was trained on *D*_train_ with deep features extracted using the zoocam_2022_04_06 model, and tested on *D*_val_ and *D*_test_ grouped together.

### Model performance metrics

To comprehensively assess classification performances among the different model architectures, we employed multiple standard metrics: accuracy, precision, recall, and F1 Score (described in details below). These metrics were necessary during both model optimization (see Model optimization strategies) and final performance comparison among optimized models (see Results).

Model optimization was based exclusively on performances measured on the validation set *D*_val_, while final performance comparison among optimized models was evaluated a single time on the held-out test set *D*_test_.

To provide insight into both overall and per-class performance, perfromance metrics were calculated in both macro-averaged and weighted-averaged forms. These metrics were computed using the scikit-learn library (Pedregosa et al., 2011), a standard in machine learning.

***Accuracy*** measures the proportion of correctly classified samples over all samples. In this work, we report both top-1 accuracy (Acc@1) and top-5 accuracy (Acc@5):

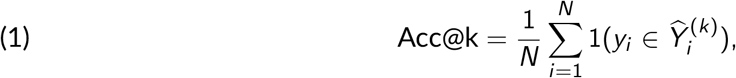

where *N* is the total number of samples, *y*_*i*_ is the ground truth label, and 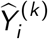 is the set of the model’s top-*k* predicted class labels for sample *i*. When *k* = 1, this reduces to standard (top-1) accuracy. 1 denotes the indicator function.

***Precision*** for a class *c* is defined as the ratio of true positives (TP) to the sum of true positives and false positives (FP):

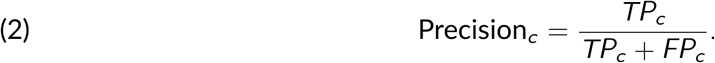

In the multiclass setting, *TP*_*c*_ denotes the number of samples correctly predicted as belonging to class *c. FP*_*c*_ counts the samples incorrectly predicted as class *c*.

***Recall*** (also called sensitivity) for class *c* measures the ratio of true positives to the sum of true positives and false negatives *FN*_*c*_, which correspond to samples of class *c* that were mistakenly classified as another class:

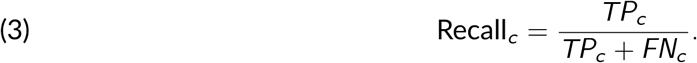

***F1 Score*** for class *c* is the harmonic mean of precision and recall:

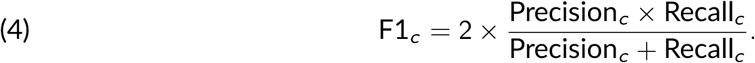

***Macro-averaged metrics*** compute the unweighted mean across classes:

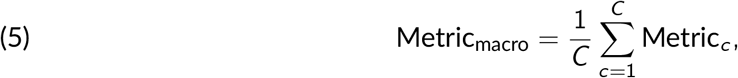

where *C* is the number of classes.

***Weighted-averaged metrics*** weight each class metric by its support (number of true samples in that class):

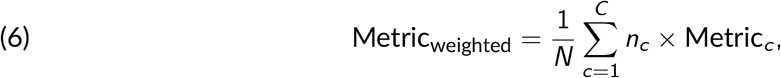

where *n*_*c*_ is the number of samples in class *c*.

In datasets with strong class imbalance such as the PlanktonFlow dataset, the choice of a method to average evaluation metrics is critical to reliably assess model performances. Metrics such as accuracy summarize performance over all samples and therefore primarily reflect performance on the most abundant classes. In our dataset, common taxa such as Daphniidae are represented by far more images than rarer taxa such as some Rotifera. Consequently, a model that performs well on these dominant taxa could achieve a high overall accuracy even if it misclassifies many samples belonging to rare taxa.

A similar limitation applies to weighted-averaged metrics (Equation 6) because each class contributes proportionally to its number of samples (*n*_*c*_ ). As a result, classes with many observations have a much larger influence on the final performance metric than rare classes. In highly imbalanced ecological datasets, this can obscure poor performance on taxa that are represented by relatively few images. To mitigate this issue, we assessed model performances using macro-averaged metrics (Equation 5), which average metrics with equal weight across classes, thus ensuring that the classifier is penalized for poor performance on rare taxa. Specifically, we selected the macro-averaged F1 Score (*F* 1_*macro*_) as our primary metric for comparing model performances.

### Model optimization strategies

All four CNN models were initialized by their developers with weights pre-trained on ImageNet, a large and generic dataset of non-planktonic images. Classically, initialization precedes a second step of model fine-tuning on a specialized dataset, an appraoch known as transfer learning from an out-of-domain source. Transfer learning has been demonstrated to be highly effective for plankton classification (Maracani et al., 2023). However, classic fine-tuning based on standard loss functions tends to favour majority classes and may fail to generalize to rare or difficult classes, which is a problem for plankton that includes many rare classes as well as classes that may overlap greatly in morphology.

To address this problem, we augmented classic fine tuning with extra optimization steps, including specialized loss functions to handle class imbalance and challenging samples, as well as hyperparameter tuning to identify optimal learning configurations (see below). Specifically, we optimized ResNet, DenseNet and EfficientNet, but we did not optimize YOLO as we chose this architecture to demonstrate the performances of an out-of-the-box deep-learning solution. EcoTaxa does not allow any model optimization by the user.

#### Loss Functions

A loss function quantifies how well the model’s predictions match the true labels. Conceptually, it can be thought of as a supervisory signal: the model attempts to minimize this loss to improve its predictions. Different loss functions emphasize different aspects of learning and can be critical in datasets characterized by class imbalance or high intra-class variability.

A common baseline is the standard cross-entropy loss *L*_CE_ (Equation 7). It treats each class equally and penalizes incorrect predictions proportionally to the predicted probability assigned to the true class:

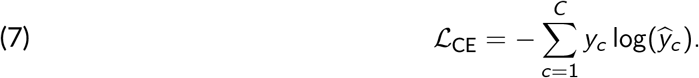

Here, *y*_*c*_ represents the ground-truth one-hot label for class *c*, and 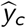 is the model’s predicted probability for that class. While effective in balanced datasets, this formulation tends to under-perform in the presence of rare classes, which are given equal treatment to frequent classes despite their limited representation.

Thus, to correct this, the weighted cross-entropy loss *L*_WCE_ (Equation 8) introduces class-specific weights *w*_*c*_ to counteract this imbalance :

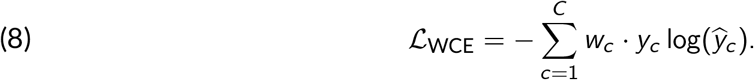

In this approach, weights are typically set inversely proportional to class frequency, effectively amplifying the contribution of rare classes during training. This ensures that the model allocates sufficient learning capacity to under-represented taxa, reducing bias toward the dominant classes.

Another approach is to use Focal Loss *L*_focal_ (Equation 9) which further improves model performance by emphasizing difficult examples (Lin et al., 2018) :

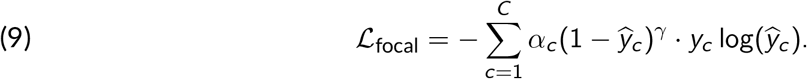

The modulating factor (1 − *ŷ*_*c*_)^*γ*^ dynamically reduces the loss contribution of well-classified “easy” examples and focuses learning on misclassified or rare instances. Here, *γ* is a focusing parameter that controls the degree of emphasis on difficult cases, while *α*_*c*_ allows optional perclass weighting. This loss function is particularly useful when both class imbalance and intra-class variability are present, as it directs learning towards the most challenging examples.

Finally, Label Smoothing (Equations 10 and 11) is a regularization technique that reduces overconfidence in predictions and encourages better generalization (Szegedy et al., 2015) :

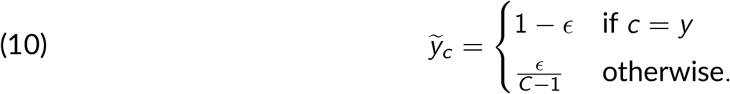

The corresponding cross-entropy loss with label smoothing becomes:

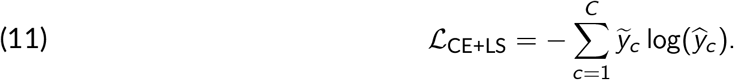

In Equation 10, the smoothing parameter *ϵ* determines the fraction of probability mass redistributed to other classes. By avoiding assigning full probability mass to a single class, label smoothing mitigates the tendency of the model to make overconfident predictions, which is particularly important when some classes are visually similar. For implementation details on considered loss functions described below, see Supplementary Information (Walter et al., 2025).

#### Hyperparameters: the knobs that control learning

Hyperparameters are parameters that define the configuration of the learning process but are not directly learned from the data. They play a critical role in determining how effectively a model can be trained and how well it generalizes to unseen data. Conceptually, hyperparameters can be thought of as ‘knobs’ that adjust how the model learns, analogous to changing the focus when examining a specimen under a microscope. Key hyperparameters include :

- Learning rate: the step size used to update the model’s parameters at each iteration, affecting convergence speed and stability.
- Batch size: the number of images processed before updating the model parameters, which influences the variance of gradient estimates and memory usage.
- Optimizer type: the algorithm used to perform parameter updates, such as Adam or AdamW, which differ in how the model adjusts weights.
- Weight decay: a regularization term that discourages overly large parameter values, helping to prevent overfitting.

Table 1 provides a detailed overview of the hyperparameters we considered during optimization, while Table 2 lists the model variants and their approximate number of parameters. Due to the large number of hyperparameters and their complex interactions, manually selecting an optimal configuration is challenging. To address this problem, we employed Bayesian optimization via Weights & Biases Sweeps (W&B, Biewald, 2020), an automated approach that efficiently explores the hyperparameter space and identifies the settings that maximize the Macro-averaged F1 Score on the validation set *D*_val_. Bayesian optimization was chosen over traditional grid or random search methods due to its efficiency in navigating large hyperparameter spaces with fewer training runs, a critical factor given the computational constraints.

**Table 1.** Hyperparameter search space explored during tuning

| Parameter | Type | Values / Range |
| --- | --- | --- |
| model_name | Categorical | {“efficientnet”, “densenet”, “resnet”} |
| model_variant | Categorical | *Model variants (see Table 2)* |
| loss_type | Categorical | {“weighted”, “focal”, “Cross-Entropy + Label Smoothing”} |
| focal_alpha | Continuous | [0.1, 0.5] |
| focal_gamma | Continuous | [1.0, 3.0] |
| use_per_class_alpha | Boolean | {true, false} |
| labelsmoothing_epsilon | Continuous | [0.05, 0.2] |
| optimizer | Categorical | {“adamw”, “adam”} |
| learning_rate | Discrete | $\{1 \times 10^{-7}, 5 \times 10^{-7}, 1 \times 10^{-6}, \dots, 2 \times 10^{-1}\}$ |
| weight_decay | Discrete | $\{1 \times 10^{-7}, 5 \times 10^{-7}, 1 \times 10^{-6}, \dots, 2 \times 10^{-1}\}$ |

**Table 2.**
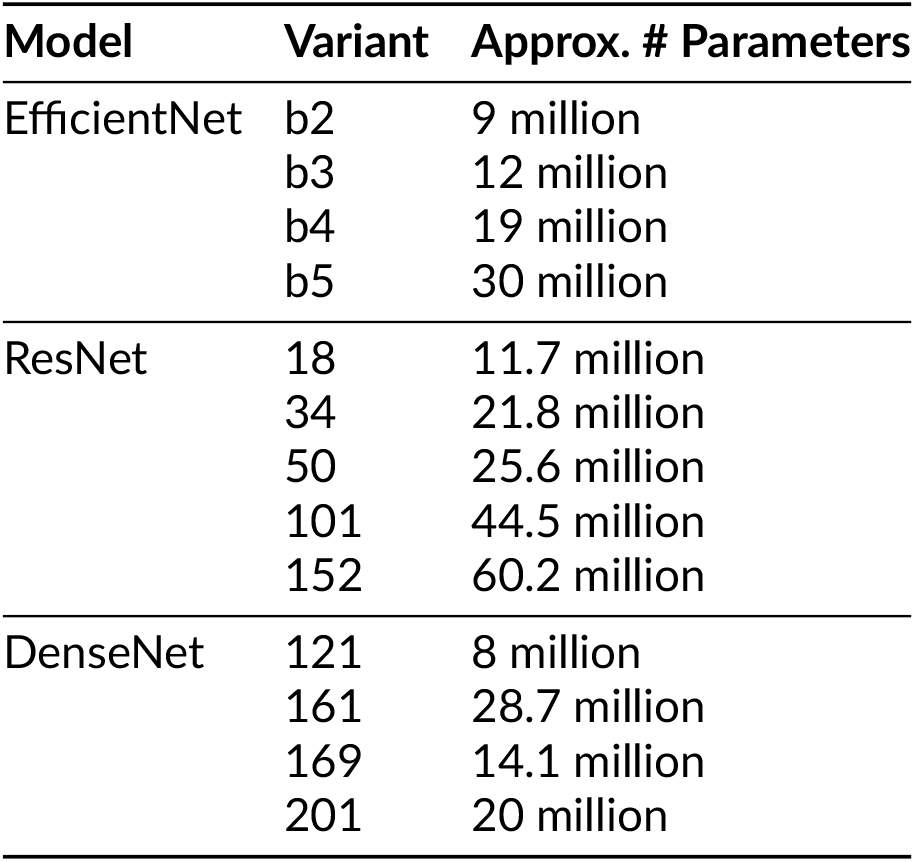
Details of model variants and their approximate number of parameters. We chose model variants that were adapted to our hardware environment (see below).

For each hyperparameter set, models were trained for a fixed number of epochs and batch size, set to 30 and 64, respectively. An epoch corresponds to a full pass of the training dataset *D*_train_ through the network, and 30 epochs were found sufficient to achieve convergence in performance while minimizing the risk of overfitting. The batch size refers to the number of training samples processed simultaneously before updating the model weights. A batch size of 64 was chosen to optimize the throughput of our hardware setup, ensuring efficient utilization of computational resources without compromising training stability. Details of the learning rate scheduler and early stopping strategy are provided in the Supplementary Information (Walter et al., 2025).

### Hardware and Software Environment

All experiments were conducted on a workstation running Ubuntu 24.04.2 LTS (Noble Numbat). The system was equipped with an NVIDIA Quadro RTX 6000 GPU featuring 24 GB of VRAM. The GPU was utilized at nearly full capacity during training, as indicated by the nvidia-smi output, with over 20 GB of memory used by the training process. The driver version used was 570.133.07, along with CUDA 12.8.

The training environment was managed via python 3.12.3 and relied heavily on GPU acceleration for faster convergence of deep learning models. The use of a professional-grade GPU ensured minimal training time, which is crucial when performing hyperparameter optimization and model evaluation across multiple configurations.

## Results

### Best-model hyperparameters

The best-performing configuration for each architecture was identified among the set of parameters that were explored by the Bayesian search (see Methods Hyperparameters). Table 3 reports the final settings retained for ResNet-152, DenseNet-161, and EfficientNet-B5. Although the three models converged toward broadly similar training regimes, subtle but meaningful differences emerged in the optimal optimization strategy. In particular, ResNet and DenseNet achieved their best performance when trained with the AdamW optimizer, whereas EfficientNet performed best with the classical Adam variant, consistent with prior reports that adaptive optimizers interact differently with depth and width scaling (Dogo et al., 2022). Likewise, EfficientNet favored a slightly larger label-smoothing parameter (*ε* = 0.12) compared to ResNet (*ε* = 0.05) and DenseNet (*ε* = 0.06), suggesting that additional regularization improved its generalization across diverse classes. The optimal learning rates and weight decay values also varied by architecture, highlighting the importance of architecture-specific tuning rather than relying on a single standardized recipe.

**Table 3.** Best hyperparameters for each model architecture on the validation set *D*_val_.

| Model | Variant | Optimizer | Loss | Learning Rate | Batch Size | Epochs | Weight Decay |
| --- | --- | --- | --- | --- | --- | --- | --- |
| ResNet | ResNet-152 | AdamW | Label Smoothing ($\epsilon=0.05$) | 1e-6 | 64 | 30 | 5e-7 |
| DenseNet | DenseNet-161 | AdamW | Label Smoothing ($\epsilon=0.06$) | 1e-6 | 64 | 30 | 1e-3 |
| EfficientNet | EfficientNet-B5 | Adam | Label Smoothing ($\epsilon=0.12$) | 1e-5 | 64 | 30 | 1e-4 |

### Relative model performances and comparison with EcoTaxa

Table 4 presents the evaluation metrics on the held-out test set *D*_test_ for each selected model alongside EcoTaxa.

**Table 4.** Performance metrics on the test set for all models. Metrics include weighted and macro-averaged recall (*R*_*w*_, *R*_*m*_), precision (*P*_*w*_, *P*_*m*_), F1 Score (*F* 1_*w*_, *F* 1_*m*_), and top-1 and top-5 accuracy (*Acc*@1, *Acc*@5). For definitions of the metrics, see Section Model performance metrics. NB.: *Acc*@5 is not available for Ecotaxa, which provides only one predicted class per image.

| Model | $R_w$ | $R_m$ | $P_w$ | $P_m$ | $F1_w$ | $F1_m$ | $Acc@1$ | $Acc@5$ |
| --- | --- | --- | --- | --- | --- | --- | --- | --- |
| ResNet | 86.15% | 84.74% | 85.76% | 84.39% | 85.77% | 84.33% | 86.53% | 98.88% |
| DenseNet | 86.59% | 84.72% | 86.24% | 84.89% | 86.27% | 84.58% | 86.40% | 98.97% |
| **EfficientNet** | 88.07% | 86.25% | 87.76% | 87.16% | 87.69% | **86.20%** | 87.90% | 98.47% |
| YOLO | 83.35% | 82.62% | 82.82% | 80.49% | 82.42% | 80.67% | 83.33% | 98.43% |
| EcoTaxa | 81.91% | 80.31% | 81.89% | 79.78% | 81.11% | 79.12% | 81.91% | NA |

ResNet, DenseNet and EfficientNet, the three models that were optimized on our dataset, stood out from the crowd. EfficientNet demonstrated the best overall performance across all metrics. In particular, EfficientNet achieved the highest macro-averaged F1 Score, suggesting a high ability to generalize across both frequent and rare classes. While EcoTaxa offered a strong traditional baseline, it lagged behind deep learning models, particularly-so in class-balanced metrics. Notably, it was surpassed by EfficientNet by a margin of +7.08 percentage points in *F* 1_*macro*_, highlighting the high potential of optimized deep learning models to automate plankton classification. To see EfficientNet’s F1 Score for all classes, see Figure S2 (Walter et al., 2025).

### Per-class analysis

While overall performance metrics provide a useful high-level summary, they often obscure important class-level behaviors—particularly in imbalanced datasets like ours. This section investigates prediction dynamics at the class level to uncover specific strengths and weaknesses of the evaluated models.

#### Class Support vs. F1 Score

To investigate whether class frequency impacted model performance, we used linear regressions of per-class F1 Scores against their corresponding support. As shown in Figure 2, regression lines were nearly flat for all models, and adjusted *R*^2^ were low, indicating almost no explanatory power. This shows that class frequency alone was not a strong predictor of classification quality, and suggests that the PlanktonFlow procedure was efficient at alleviating the classification problems that arise from class imbalance (a formal test of this hypothesis would require to run the analysis without the pre-processing procedure).

**Figure 2.**
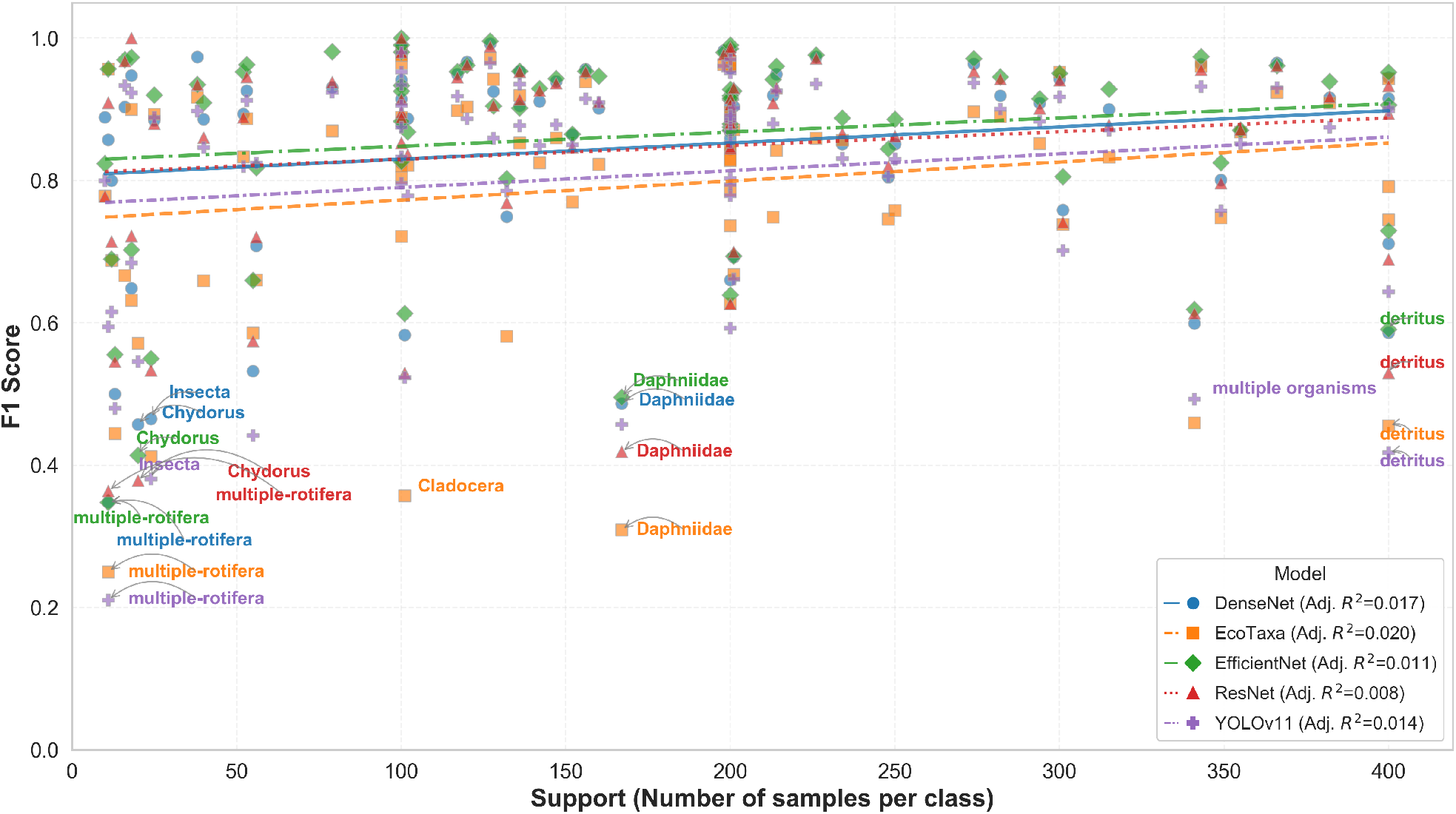
Scatter plot of per-class F1 Score versus class support in the test set *D*_test_ for each model. Rare classes generally perform worse, but some achieve strong F1 Scores.

#### Worst Performing Classes

To better understand model weaknesses, we analysed the ten lowest-performing classes per model in terms of F1 Score. Poor performance was often observed in classes with high intra-class variability (e.g., “detritus”, “multiple organisms”, “Insecta”), or among overlapping classes like “Cladocera” and “damaged_Cladocera” (Figure 3). Nested taxonomic groups were also difficult to classify due to their inherent morphological similarity (e.g., both “Daphniidae” and “Ceriodaphnia” are taxonomically nested in “Cladocera”, Figure 3). While the specific failing classes differ across models, some categories appear recurrently among the bottom ten, suggesting inherent classification difficulty (e.g., multiple-rotifera).

**Figure 3.**
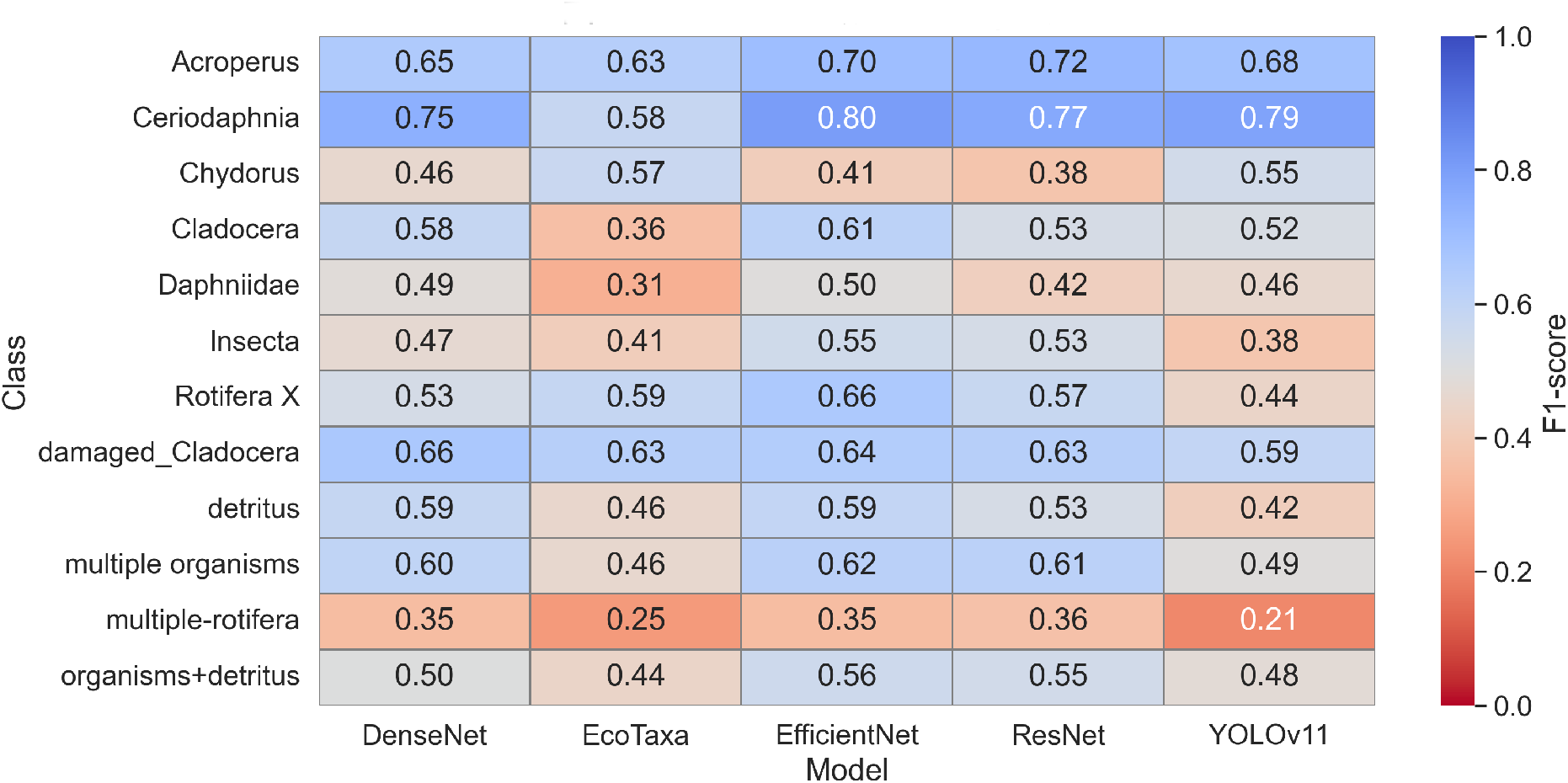
Per-model F1 Scores for the 10 worst-performing classes.

#### Top confused class pairs

To uncover systematic sources of misclassification, we identified the five most confused class pairs for each model by examining off-diagonal elements in the confusion matrices (Figure 4). Again, we observed that top most confused class pairs are classes with high intra-class variability, with the leading pair being “detritus-dark” confused for “detritus”. Again, confusion was also due to “taxonomic nestedness”, where one class was confused for a higher class in the taxonomy (e.g., “Cyclopoida” confused for “Copepoda”, Figure 4). Finally, we *a posteriori* found that confusion among the “Chydoridae” and “Ostracoda” classes were due mainly to human errors in the dataset. After dataset cleaning for these two particular classes and re-running the pipeline, the total number of confused images dropped from 50 to 15.

**Figure 4.**
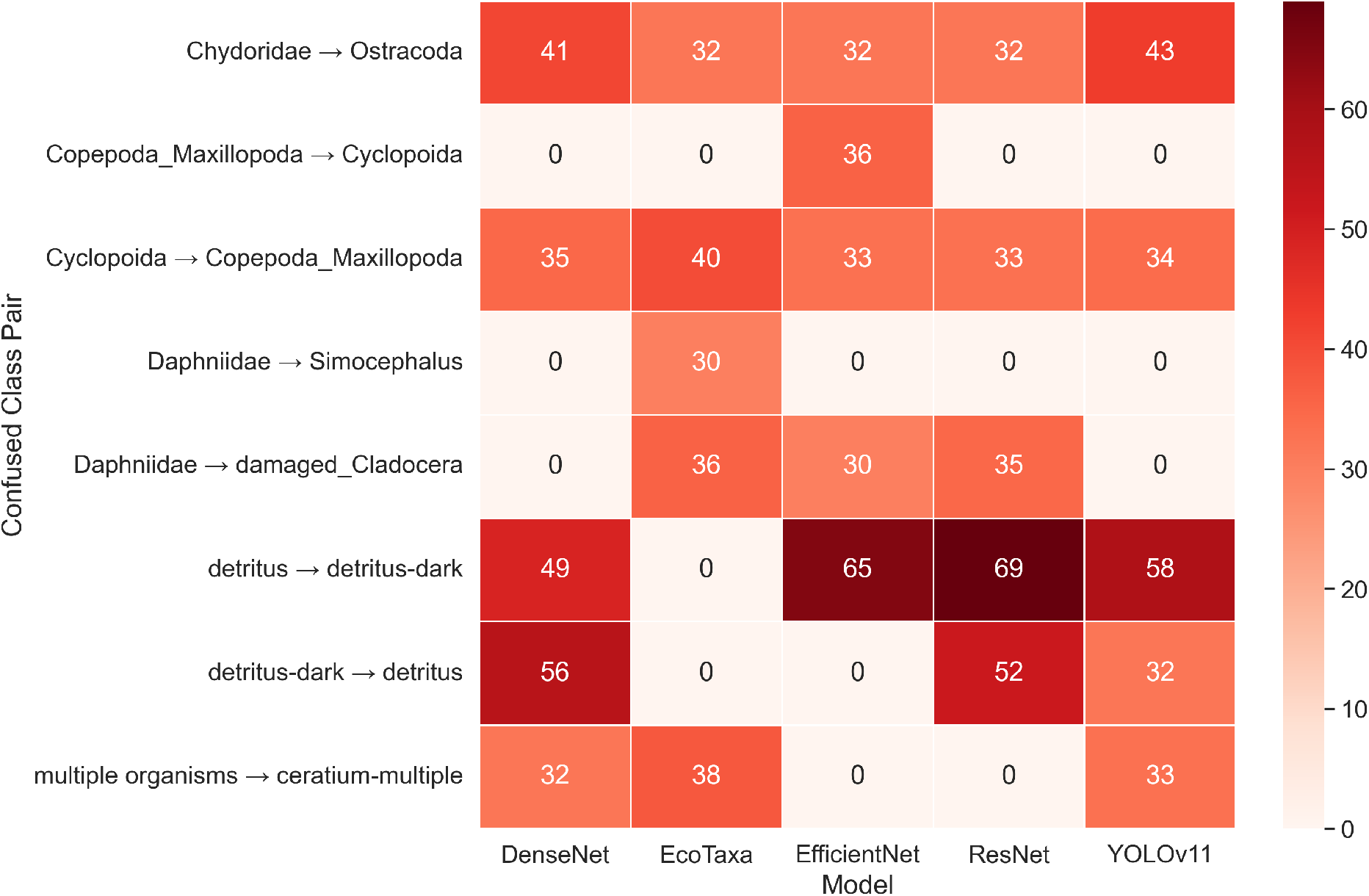
Top 5 most confused class pairs per model based on confusion matrix analysis. A→B indicates that A class was confused for class B. Numbers indicate the number of confused images.

#### Error distribution among difficult classes (EfficientNet)

Given that EfficientNet was the best performing model overall, we performed a fine-grained error analysis focused on this architecture. Figure 5 presents the 20 classes with the highest total number of errors, computed as the sum of false positives and false negatives. The distribution of error types across these classes appeared relatively balanced, with false positives and false negatives contributing in roughly equal proportions. This suggests that EfficientNet did not exhibit a strong bias toward over- or under-predicting specific classes among its most error-prone predictions, but rather that the model struggled equally with both types of misclassification in these challenging classes.

**Figure 5.**
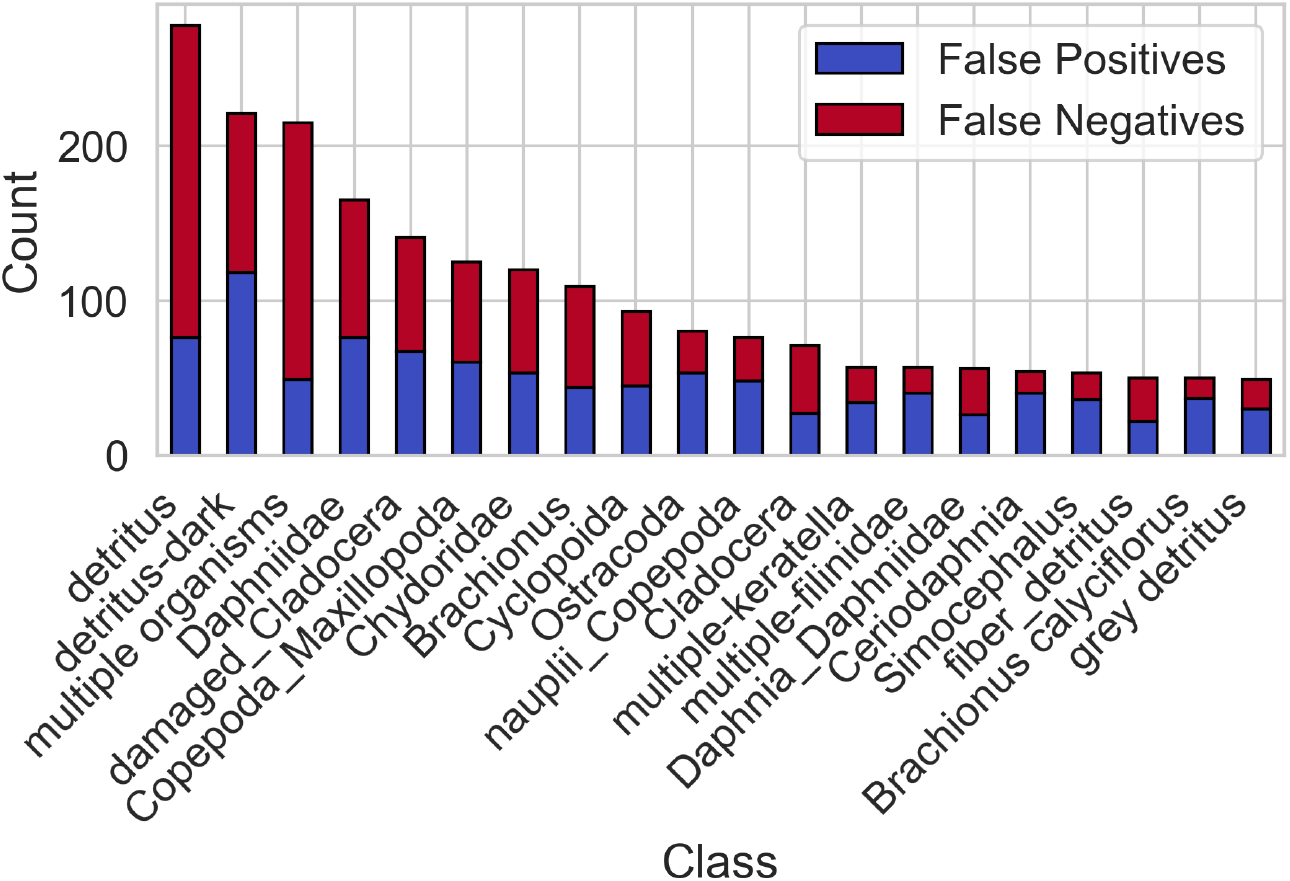
Top 20 classes by total error count for EfficientNet, with separate bars for false positives and false negatives.

## Discussion

Our results demonstrated the capabilities of PlanktonFlow for the implementation of a complete deep-learning pipeline, including a performance comparison among multiple models (for a hands-on tutorial on how to reproduce the results, see the next section). On our custom dataset, the best overall performance was achieved by EfficientNet-B5, while both ResNet and DenseNet achieved comparable results within a 1-2% margin. YOLO was lagging behind, illustrating the limits of out-of-the-box solutions when they are not optimized on the dataset at hand. Finally, our results show that EcoTaxa delivered quite decent performances, but was neatly surpassed by optimized CNNs. The suitability of EfficientNets for plankton classification was already highlighted (Kyathanahally et al., 2021), suggesting that their performance could possibly be relatively robust to dataset shifts (Chen et al., 2025). However, we recommend PlanktonFlow users to systematically compare the performances of multiple optimized CNNs on their own dataset so as to choose the best-performing one. PlanktonFlow was specifically designed to ease such model comparison.

Our study further illustrates the already-known power of CNNs for taxonomic classification of plankton images (e.g., Kyathanahally et al., 2021, Kyathanahally et al., 2022, Eerola et al., 2024, Zheng et al., 2026). Our best model, EfficientNetB5, achieved a 86% Macro-averaged F1 Score, which is close the ∼ 90% macro-averaged F1 Scores typically achieved by state-of-the-art CNNs trained on benchmark datasets (Kyathanahally et al., 2021, Kyathanahally et al., 2022, Zheng et al., 2026). This is remarquable given that our study included the difficult classes that are typical of non-benchmark, real world datasets (diverse detritus classes, multiple objects on same image, overlapping classes, or nested taxonomic classes), and also given that the dataset included annotation errors among two classes. From a biological perspective, overlapping and taxonomically-nested classes are not necessarily problematic because they share some taxonomic information and they can thus be merged at the post-processing stage (albeit at the cost of a lower taxonomic resolution). In contrast, annotations errors, if they were not a problem for our purpose of demonstrating the usage of PlanktonFlow, should systematically be tracked and eliminated before any final analysis.

### PlanktonFlow : a hands-on tutorial

The whole pipeline is downloadable from GitHub (PlanktonFlow) and comes with a full-fledged documentation.

#### Typical workflow with the pipeline

PlanktonFlow is entirely configuration-driven using dedicated YAML configuration files, in which detailed explanations are provided in the form of comments. Using these comments, users are expected to adapt these files to their own dataset and experimental design. This ensures full reproducibility while keeping commands themselves simple. A typical workflow involves a sequence of preprocessing, training, and inference. Each of these three operations has its own YAML configuration file and may be performed independently.

#### Data preprocessing

PlanktonFlow accommodates three common data formats:

1. **Hierarchical dataset**: classical folder-based structure with one folder per image class.
2. **Simple CSV/TSV mapping**: one single folder containing all images mixed together, accompanied by a basic, user-generated two-column CSV/TSV file simply mapping each image filename to its class label.
3. **EcoTaxa export**: a specialized parser designed to directly ingest the complex, multi-column TSV archives exported by the EcoTaxa web platform. It automatically navigates the EcoTaxa-specific metadata structure (e.g., extracting labels from the object_annota tion_category column) to associate classes with images, eliminating the need for manual file reformatting by the user.

Users can customize preprocessing through configuration files to:

- Prune classes based on a minimum and maximum number of images.
- Perform data augmentation to balance the dataset.
- Optionally remove scale bars using a pre-trained YOLOv8 model.
- Automatically split data into training, validation, and test sets.

At the end of this step, the dataset should be ready to train a model with.

#### Training

Training is fully configurable via YAML files. Users can select a model architecture, model variants, loss functions (e.g., focal loss, label smoothing, weighted cross-entropy), optimizers etc. Metrics are tracked in real-time, either locally or online through Weights & Biases (W&B). Pre-configured hyperparameters have been validated on our plankton dataset, allowing FlowCAM users to achieve good performance out-of-the-box. However, PlanktonFlow also makes it possible to optimize hyperparameters on your own dataset.

#### Hyperparameter optimization

Hyperparameter optimization in PlanktonFlow operates W&B, which provides an efficient and reproducible framework for exploring large hyperparameter spaces. Instead of requiring the user to manually test parameters such as learning rate, weight decay, batch size, or optimizer type, the pipeline automatically launches a series of training runs with different configurations sampled from a predefined search space (specified in the YAML sweep configuration file). The underlying algorithm in W&B prioritizes promising regions of the search space, thereby reducing the number of runs needed compared to a naive grid search.

From the PlanktonFlow user’s perspective, initiating a sweep requires only pointing to the provided sweep configuration file. The sweep will continue to launch successive training jobs until it is manually stopped. Typically, users are advised to monitor validation metrics in the W&B dashboard and stop the sweep once these metrics show signs of stable convergence across runs. At this point, the best-performing hyperparameter set can be automatically exported into a dedicated YAML configuration file for further training or inference. PlanktonFlow also saves every trained model together with its configuration, ensuring that optimal runs are reproducible and can be reused in subsequent analyses.

*Inference*. PlanktonFlow allows batch inference on new image directories. Users can:

- Retrieve top-K predictions per image.
- Apply optional preprocessing (e.g., scale bar removal, see above).
- Export results to CSV for further analysis.

Device-aware execution ensures models run on CPU or GPU depending on availability. Inference outputs include detailed prediction probabilities and optional preprocessing steps applied automatically.

#### Monitoring and results analysis

If W&B is not used, a custom logging module records training and evaluation metrics in real time. Metrics are stored in CSV files, enabling visualization of training curves, confusion matrices, F1 Scores, and class-wise performance. An accompanying Jupyter notebook (results_analysis.ipynb) facilitates comprehensive evaluation and figure generation.

#### PlanktonFlow usage

After installation of PlanktonFlow following the guidelines in the readme file, and before executing the pipeline, users must download the published dataset, or provide their own data in the correct directory. Then, a complete workflow can be executed as follows:

1. Preprocess data:

~~~
python 3 run_preprocessing . py -- config configs/ preprocessing /
    PreprocessWith Hierarchical . yaml
~~~
2. Train a model:

~~~
python 3 run_training . py -- config configs/ training /
  Train DefaultEfficientNet . yaml
~~~
3. Make predictions:

~~~
python 3 run_inference . py -- config configs/ inference / DefaultInference .
  yaml
~~~
4. Optimize:

~~~
python 3 run_sweep . py -- sweep_config configs/ sweeps/ densenet_sweep .
  yaml
~~~

## Conclusion

A major strength of PlanktonFlow is its high degree of automation and scalability: all core stages, from data preprocessing to training and inference, are fully automated, minimizing the need for human intervention and making the approach highly suitable for large-scale or continuous monitoring workflows. Also, although the models were trained on a specific plankton dataset, the underlying architecture and modular structure of the pipeline promote generalizability. With only minimal changes to the input format or preprocessing configuration, PlanktonFlow could be adapted to other image-acquisition devices or taxonomic groups. Our hope is that the community will use and further develop PlanktonFlow. We foresee that future work could leverage Vision transformers, semi-supervised learning, and adapt the pipeline to other imaging and biological systems.

## Acknowledgements

The authors thank the Villefranche sur Mer Quantitative Imaging Platform (PIQv) for supporting the community of plankton biologists with free and high-quality services. In particular, our research in general, and this study in particular, rely heavily on ZooProcess and EcoTaxa.

## Data, script, code, and supplementary information availability

Datasets are available online at https://doi.org/10.5281/zenodo.16840846 (Gorzerino et al., 2025). Scripts and codes for PlanktonFlow are available at https://github.com/ziraax/PlanktonFlow (Walter, 2025). Supplementary information for figure reproduction and implementation details is available at https://doi.org/10.5281/zenodo.19222299 (Walter et al., 2025).

## Supplementary Information

### S1 Supplementary Methods

#### S1.1 Loss function algorithms

##### Algorithm 1

Computation of Focal Loss (per sample)

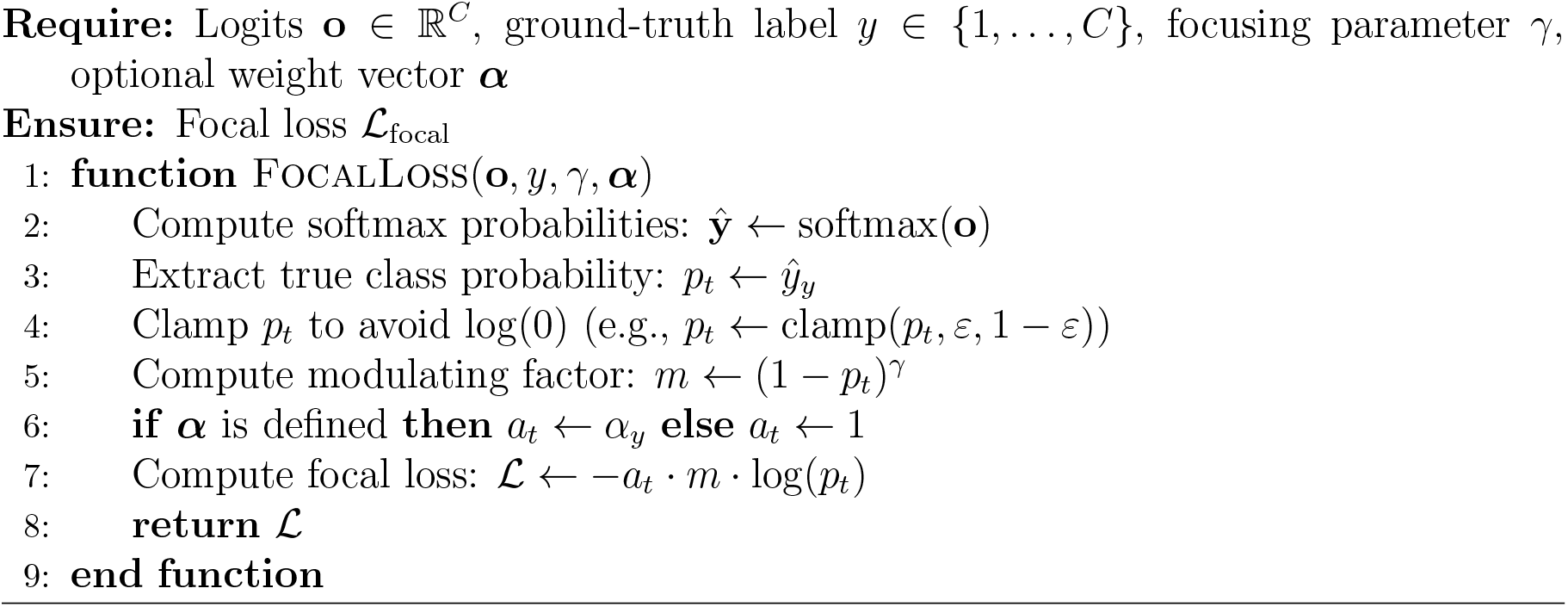

##### Algorithm 2

Computation of Label Smoothing Loss (redistributing smoothing over incorrect classes)

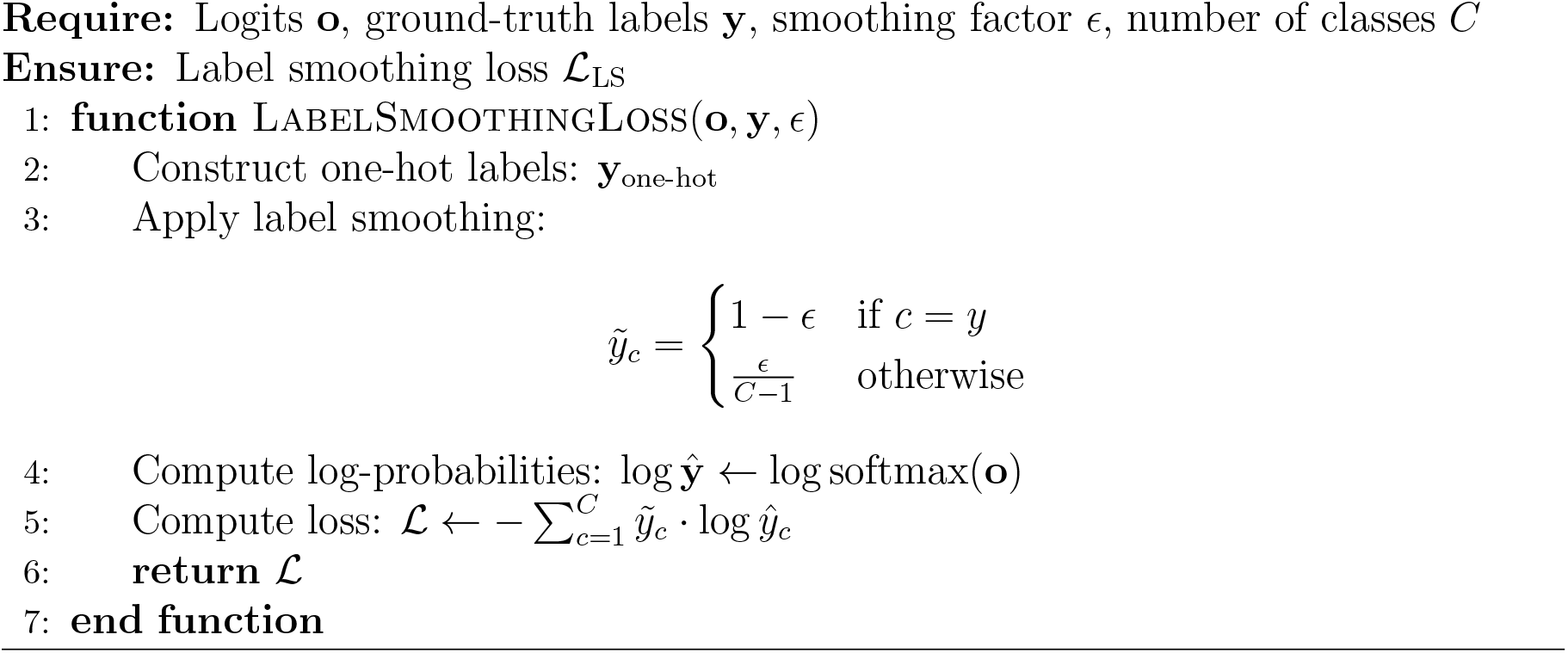

#### S1.2 Learning Rate Scheduling

In addition to optimizer selection, the learning rate was dynamically adjusted throughout training using a cosine annealing schedule. Specifically, we used PyTorch’s CosineAnnealingLR, which gradually decreases the learning rate following a cosine function over a fixed number of epochs. This approach encourages larger updates at the beginning of training and finer adjustments toward the end, helping the model settle into better minima.

At each epoch *t*, the learning rate *η*_*t*_ is computed as:

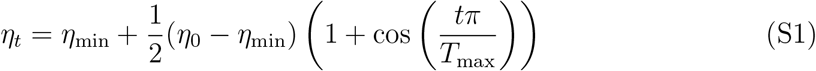

where *η*_0_ is the initial learning rate, *η*_min_ is the minimum learning rate (often set to 0), *t* is the current epoch, and *T*_max_ is the total number of epochs before a full cosine cycle completes.

#### S1.3 Early Stopping Strategy

To prevent unnecessary training time, an **early stopping** mechanism was implemented (see Algorithm 3). This method monitors the validation loss and, optionally, the validation accuracy, halting training when no improvement is observed for a specified number of consecutive epochs (referred to as the *patience* parameter).

Our implementation allows for two simultaneous criteria:

- Validation loss must improve by at least a threshold *δ* (default: 0).
- If enabled, validation accuracy must also improve by at least *δ*.

Formally, let *L*_*t*_ and *A*_*t*_ denote the validation loss and accuracy at epoch *t*. Training stops if:

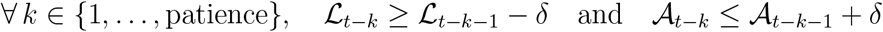

This dual-metric early stopping criterion improves robustness over single-metric strategies, particularly in settings where accuracy and loss are not perfectly aligned.

##### Algorithm 3

Early Stopping Procedure

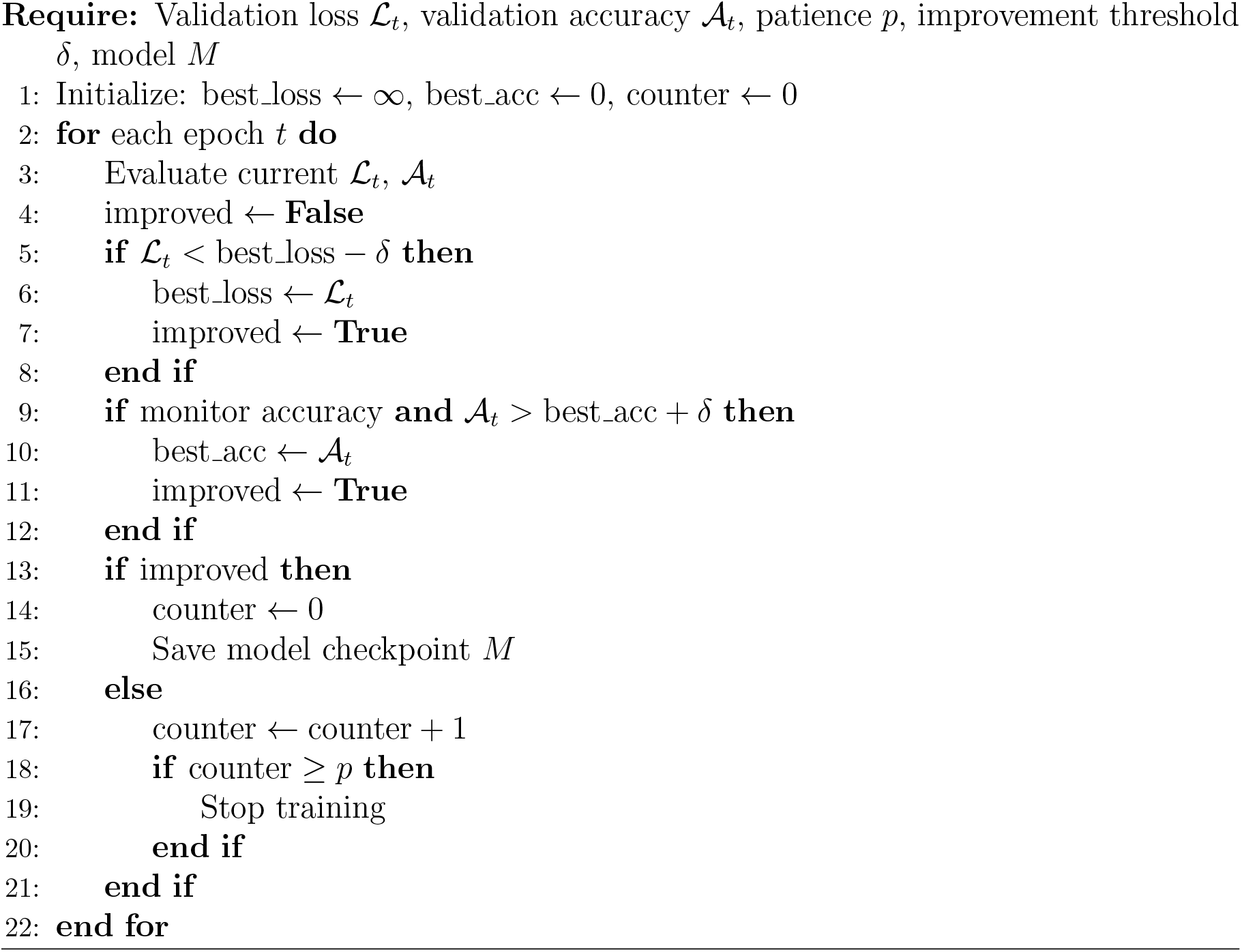

### S2 Additional Figures

**Figure S1:**
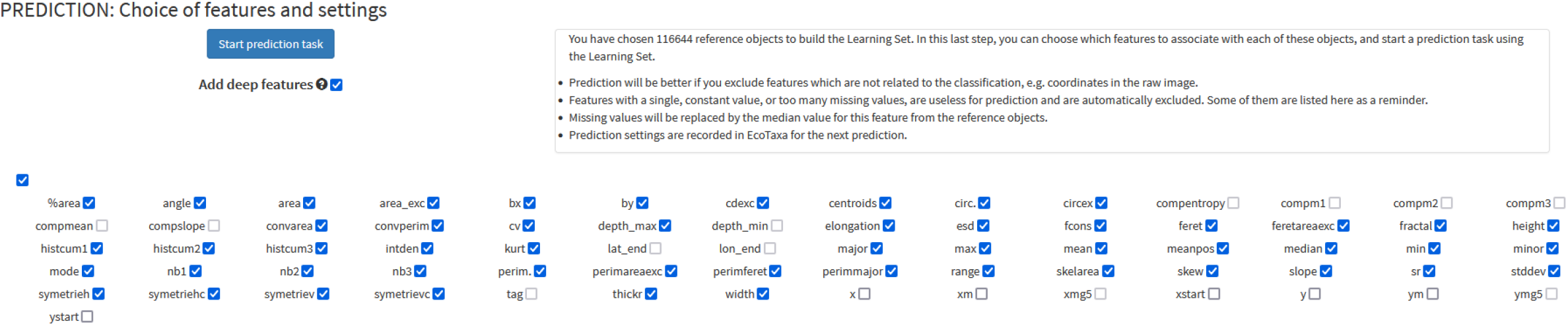
EcoTaxa selected parameters.

**Figure S2:**
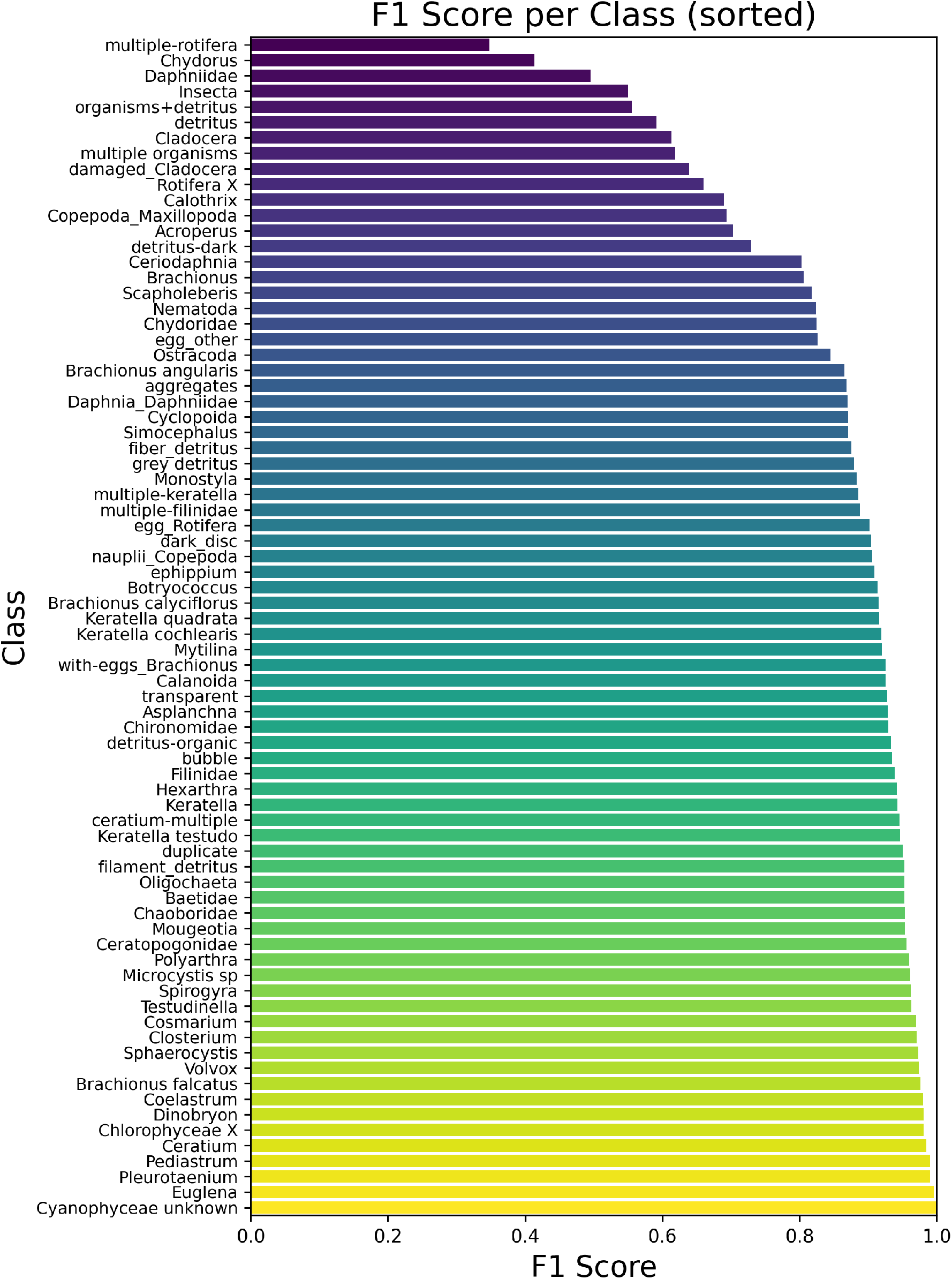
F1 Score per class for the best trained model (EfficientNet-B5)

## Footnotes

1 https://github.com/ultralytics/ultralytics

## References

Besson M, Alison J, Bjerge K, Gorochowski TE, Høye TT, Jucker T, Mann HMR, Clements CF (2022). Towards the fully automated monitoring of ecological communities. Ecology Letters 25, 2753–2775. 10.1111/ele.14123.

Biewald L (2020). Experiment tracking with Weights and Biases. Software available from wandb.com. URL: https://www.wandb.com/.

Buslaev A, Parinov E, Khvedchenya VI, Iglovikov, Kalinin AA (2018). Albumentations: fast and flexible image augmentations. ArXiv e-prints. eprint: 1809.06839.

Chen C, Kyathanahally SP, Reyes M, Merkli S, Merz E, Francazi E, Hoege M, Pomati F, Baity-Jesi M (2025). Producing plankton classifiers that are robust to dataset shift. Limnology and Oceanography: Methods 23, 39–66. 10.1002/lom3.10659. URL: https://aslopubs.onlinelibrary.wiley.com/doi/abs/10.1002/lom3.10659.

Dézerald O, Gorzerino C, Fourcy D, Collinet M, Coudreuse J, Benneveault Y, Marchand F, Azam D, Edeline E (2023). Eco-evolutionary effects of invasive crayfish on aquatic ecosystems. In: Symposium AQUACOSM+. Antalya (Turkey), Turkey. URL: https://hal.science/hal-05483521.

Dogo EM, Afolabi OJ, Twala B (2022). On the relative impact of optimizers on convolutional neural networks with varying depth and width for image classification. Applied Sciences 12. 10.3390/app122311976.

Eerola T, Batrakhanov D, Barazandeh NV, et al. (2024). Survey of automatic plankton image recognition: challenges, existing solutions and future perspectives. Artificial Intelligence Review 57, 114. 10.1007/s10462-024-10745-y.

Faillettaz R, Picheral M, Luo JY, Guigand C, Cowen RK, Irisson JO (2016). Imperfect automatic image classification successfully describes plankton distribution patterns. Computer Vision in Oceanography 15-16, 60–77. 10.1016/j.mio.2016.04.003. URL: https://www.sciencedirect.com/science/article/pii/S2211122015300177.

Gorzerino C, Walter H, Edeline E, Martignac F (2025). PlanktonFlow76 - A curated FlowCam dataset for plankton classification. Zenodo. 10.5281/zenodo.16840846.

He K, Zhang X, Ren S, Sun J (2016). Deep residual learning for image recognition. In: 2016 IEEE conference on computer vision and pattern recognition (CVPR), pp. 770–778. 10.1109/CVPR.2016.90.

Huang G, Liu Z, van der Maaten L, Weinberger KQ (2018). Densely connected convolutional networks. arXiv: 1608.06993 [cs.CV]. URL: https://arxiv.org/abs/1608.06993.

Irisson JO, Salinas L, Colin S, Complex T, Picheral M (2022). EcoTaxa: a tool to support the taxonomic classification of large datasets through supervised machine learning. In: SFEcologie 2022. Metz, France. URL: https://hal.science/hal-04026447.

Jalabert L, Elineau A, Brandão M, Picheral M (2025). Zooscan-Zooprocess user manual and procedures at the Quantitative Imaging Platform of Villefranchesur-Mer (PIQv). 10.5281/zenodo.15114134.

Khanam R, Hussain M (2024). YOLOv11: an overview of the key architectural enhancements. arXiv: 2410.17725 [cs.CV]. URL: https://arxiv.org/abs/2410.17725.

Kraft K, Velhonoja O, Eerola T, Suikkanen S, Tamminen T, Haraguchi L, Ylöstalo P, Kielosto S, Johansson M, Lensu L, Kälviäinen H, Haario H, Seppälä J (2022). Towards operational phytoplankton recognition with automated high-throughput imaging, near-real-time data processing, and convolutional neural networks. Frontiers in Marine Science Volume 9 - 2022. 10.3389/fmars.2022.867695. URL: https://www.frontiersin.org/journals/marine-science/articles/10.3389/fmars.2022.867695.

Kyathanahally SP, Hardeman T, Reyes M, Merz E, Bulas T, Brun P, Pomati F, Baity-Jesi M (2022). Ensembles of data-efficient vision transformers as a new paradigm for automated classification in ecology. Scientific Reports 12, 18590. 10.1038/s41598-022-21910-0. URL: https://doi.org/10.1038/s41598-022-21910-0.

Kyathanahally SP, Hardeman T, Merz E, Bulas T, Reyes M, Isles P, Pomati F, Baity-Jesi M (2021). Deep learning classification of lake zooplankton. Frontiers in Microbiology Volume 12-2021. 10.3389/fmicb.2021.746297. URL: https://www.frontiersin.org/journals/microbiology/articles/10.3389/fmicb.2021.746297.

Lin TY, Goyal P, Girshick R, He K, Dollár P (2018). Focal loss for dense object detection. arXiv 1708.02002 [cs.CV]. URL: https://arxiv.org/abs/1708.02002.

Lumini A, Nanni L (2019). Deep learning and transfer learning features for plankton classification. Ecological Informatics 51, 33–43. 10.1016/j.ecoinf.2019.02.007. URL: https://www.sciencedirect.com/science/article/pii/S1574954118303054.

Maracani A, Pastore VP, Natale L, et al. (2023). In-domain versus out-of-domain transfer learning in plankton image classification. Scientific Reports 13, 10443. 10.1038/s41598-023-37627-7.

Merz E, Kozakiewicz T, Reyes M, Ebi C, Isles P, Baity-Jesi M, Roberts P, Jaffe JS, Dennis SR, Hardeman T, Stevens N, Lorimer T, Pomati F (2021). Underwater dual-magnification imaging for automated lake plankton monitoring. Water Research 203, 117524. 10.1016/j.watres.2021.117524. URL: https://www.sciencedirect.com/science/article/pii/S004313542100720X.

Olson RJ, Sosik HM (2007). A submersible imaging-in-flow instrument to analyze nano-and microplankton: imaging FlowCytobot. Limnology and Oceanography: Methods 5, 195–203. 10.4319/lom.2007.5.195.

Orenstein EC, Beijbom O, Peacock EE, Sosik HM (2015). WHOI-Plankton-A large scale fine grained visual recognition benchmark dataset for plankton classification. arXiv: 1510.00745 [cs.CV]. URL: https://arxiv.org/abs/1510.00745.

Owen BM, Hallett CS, Cosgrove JJ, Tweedley JR, Moheimani NR (2022). Reporting of methods for automated devices: a systematic review and recommendation for studies using FlowCam for phytoplankton. Limnology and Oceanography: Methods 20, 400–427. 10.1002/lom3.10496.

Pedregosa F, Varoquaux G, Gramfort A, Michel V, Thirion B, Grisel O, Blondel M, Prettenhofer P, Weiss R, Dubourg V, Vanderplas J, Passos A, Cournapeau D, Brucher M, Perrot M, Duchesnay E (2011). Scikit-learn: machine learning in Python. Journal of Machine Learning Research 12, 2825–2830.

Picheral M, Colin S, Irisson JO (2025). EcoTaxa, a tool for the taxonomic classification of images. http://ecotaxa.obs-vlfr.fr. Accessed: 2025-05-29.

Picheral M, Guidi L, Stemmann L, Karl DM, Iddaoud G, Gorsky G (2010). The Underwater Vision Profiler 5: an advanced instrument for high spatial resolution studies of particle size spectra and zooplankton. Limnology and Oceanography: Methods 8, 462–473. 10.4319/lom.2010.8.462.

Pollina T, Larson AG, Lombard F, Li H, Le Guen D, Colin S, de Vargas C, Prakash M (2022). PlanktoScope: affordable modular quantitative imaging platform for citizen oceanography. Frontiers in Marine Science Volume 9 - 2022. URL: https://www.frontiersin.org/journals/marine-science/articles/10.3389/fmars.2022.949428.

Sieracki C, Sieracki M, Yentsch C (1998). An imaging-in-flow system for automated analysis of marine microplankton. Marine Ecology-progress Series 168, 285–296. 10.3354/meps168285.

Szegedy C, Vanhoucke V, Ioffe S, Shlens J, Wojna Z (2015). Rethinking the inception architecture for computer vision. arXiv: 1512.00567 [cs.CV]. URL: https://arxiv.org/abs/1512.00567.

Tan M, Le QV (2020). EfficientNet: rethinking model scaling for convolutional neural networks. arXiv: 1905.11946 [cs.LG]. URL: https://arxiv.org/abs/1905.11946.

Walter H (2025). Deep learning classification pipeline for automatic plankton classification. Version 1.0.0. URL: https://github.com/ziraax/PlanktonFlow.git.

Walter H, Gorzerino C, Collinet M, Porcon B, Martignac F, Edeline E (2025). PlanktonFlow : supplementary information & Best model weight. 10.5281/zenodo.19222299.

Weinstein BG (2018). A computer vision for animal ecology. Journal of Animal Ecology 87, 533–545. 10.1111/1365-2656.12780.

Zheng J, Sun Z, Guo R, Wang R, Jin D, Hu J, Xing X, Tong M, Wang P (2026). High-throughput and rapid classification on harmful algal bloom species based on mega image database and artificial intelligence. Marine Pollution Bulletin 224, 119170. 10.1016/j.marpolbul.2025.119170. URL: https://www.sciencedirect.com/science/article/pii/S0025326X25016467.

